# Gut-testis axis: microbiota-(n-3) PUFA improving semen quality in type 1 diabetes

**DOI:** 10.1101/2021.11.24.469971

**Authors:** Yanan Hao, Yanni Feng, Xiaowei Yan, Liang Chen, Ruqing Zhong, Xiangfang Tang, Wei Shen, Qingyuan Sun, Zhongyi Sun, Yonglin Ren, Hongfu Zhang, Yong Zhao

## Abstract

Gut dysbiosis and type 1 diabetes (T1D) are closely related, and gut dysbiosis and male infertility are correlated, too. Moreover, most male T1D patients are of active reproductive age. Therefore, it is crucial to explore possible means for improving their semen quality. Here, we found that fecal microbiota transplantation (FMT) from alginate oligosaccharide (AOS) improved gut microbiota (A10-FMT) significantly decreased blood glucose and glycogen, and increased semen quality in streptozotocin-induced T1D subjects. A10-FMT improved T1D-disturbed gut microbiota, especially the increase in small intestinal *lactobacillus*, and blood and testicular metabolome to produce n-3 polyunsaturated fatty acid (PUFA) docosahexaenoic acid (DHA) and eicosapentaenoic acid (EPA) to ameliorate spermatogenesis and semen quality. Moreover, A10-FMT can improve spleen and liver function to strengthen the systemic environment for sperm development. FMT from gut microbiota of control animals (Con-FMT) produced some beneficial effects; however, to a smaller extent. Thus, AOS improved gut microbiota may be a useful protocol for improving semen quality and male fertility in T1D patients.

**Importance:** Clinical data suggest that male reproductive dysfunction especially infertility is a critical issue for type 1 diabetic patient (T1D) because most of them are at the reproductive age. Gut dysbiosis is involved in T1D related male infertility. However, improved gut microbiota can be used to improve spermatogenesis and male fertility in T1D remains incompletely understood. We discovered that alginate oligosaccharide-improved gut microbiota (A10-FMT) significantly ameliorated spermatogenesis and semen quality. AOS-improved gut microbiota (specific microbes) may serve as a novel, promising therapeutic approach for the improvement of semen quality and male fertility in T1D patients.

## Introduction

Type 1 diabetes (T1D), one of the most common metabolic disorders in children and young adults, is a multifactorial, immune-mediated disease that is characterized by the progressive destruction of autologous insulin-producing beta cells in the pancreas, and an increase in blood glucose levels (hyperglycemia)^1–3^. Diabetic hyperglycemia leads to further disorder, including cardiovascular disease, neuropathy, nephropathy, retinopathy, and male impotence. Moreover, clinical data from males with hyperglycemia-induced reproductive dysfunction are reported in most T1D studies^4^. This increase in diabetes in young persons is of great concern as it will increase fertility related disorders during their reproductive lifespan^5^. Furthermore, the frequency of diabetes mellitus (DM) in males is higher than in females, and the incidence of infertility among diabetic males is more common, which will likely contribute to the reduction of global birth rates and fertility^4^. Many investigations have found DM-induced male infertility at multiple levels, such as changes in spermatogenesis, testes, ejaculatory function and libido^5, 6^ Additionally, cross-sectional studies reported that at the time of infertility diagnosis, young men are already less healthy than their fertile peers which suggests that male reproductive and somatic health are tightly corelated^7^.

Although T1D has a strong genetic basis, epigenetic and environmental factors (hygiene, antibiotic use, and diet) are involved in its development^2, 8^. Furthermore, all of these potential environmental risk factors are related to the intestine and its microbiota. Specific alterations in the diversity of intestinal microflora have been reported to be one characteristic of diabetic patients by the epidemiological investigations^9–11^. Interaction of the gut microbes with the innate immune system plays vital roles in the development of T1D^8^. It has been reported that a reduction in bacterial and functional diversity, and community stability are found in preclinical T1D patients, on the other hand *Bacteroidetes* is dominated^12–15^. The presence of small intestinal *Prevotella* are known to be inversely related to pancreatic beta cell function, however small intestinal *Desulfovibrio* are involved in preserved beta cell function^8^.

Very recently, it has been established that the gut microbiota plays crucial roles in spermatogenesis and male fertility^16–19^. Studies demonstrate a strong link between testicular function and the regulation of gut microbiota via host metabolomes, as beneficial microbiota have been shown to significantly improve busulfan impaired spermatogenesis and semen quality^17, 18^. Since noting the high incidence of infertility in T1D male patients, many investigators have tried to improve the semen quality and fertility in T1D induced animal models^20, 21^. It is reported that resveratrol attenuates reproductive alterations in T1D-induced rats with improvements in glycemic level, sperm quantitative and qualitative parameters, and the hormonal profile^20^. Liu et al., found that metformin ameliorates testicular damage (90% of male DM patients have varying degrees of testicular dysfunction) in male mice with streptozotocin (STZ)-induced T1D through the PK2/PKR Pathway^21^. However, it is unknown whether impaired spermatogenesis and semen quality in the T1D condition can be improved by FMT or specific microbes. Therefore, this study aimed to explore possible improvements in spermatogenesis and semen quality made by alginate oligosaccharide (AOS) benefited microbiota which have previously been found to ameliorate busulfan or high fat diet disrupted spermatogenesis and male fertility.

## Materials and Methods

### Study design

All animal procedures used in this study were approved by the Animal Care and Use Committee of the Institute of Animal Sciences of Chinese Academy of Agricultural Sciences (IAS2020-106). Mice were maintained in specific pathogen-free (SPF) environment under a light: dark cycle of 12:12 h, at a temperature of 23 ℃ and humidity of 50%–70%; they had free access to food (chow diet) and water^17, 18, 22^.

### Experiment I: Mouse small intestine microbiota collection^17, 18^

Three-week-old ICR male mice were dosed with ddH_2_O as the control or AOS 10 mg/kg BW via oral gavage (0.1 ml/mouse/d). AOS dosing solution was freshly prepared on a daily basis and delivered every morning for three weeks. There were two groups (30 mice/treatment): (1) Control (ddH_2_O); (2) A10 (AOS 10 mg/kg BW). After three weeks treatment, the animals were maintained on regular diet for three more days (no treatment). Then the mice were humanely euthanized to collect small intestinal luminal content (microbiota).

### Experiment II: STZ treatment and microbiota transplants (FMT)^17, 18, 23, 24^

The small intestine luminal content (microbiota) from each group was pooled and homogenized, diluted 1:1 in 20% sterile glycerol (saline) and frozen. Before inoculation, fecal samples were diluted in sterile saline to a working concentration of 0.05 g/ml and filtered through a 70-μm cell strainer. Five-week-old ICR male mice were used in current investigation. There were four treatment groups (30 mice/treatment): (1) Control (Regular diet plus Saline); (2) STZ (One dose STZ at 135mg/kg body weight after preliminary screening) ^20^; (3) Con-FMT [STZ plus gut microbiota from control mice (Experiment I)]; (4) A10-FMT [STZ plus gut microbiota from AOS 10 mg/kg mice (Experiment I)]. STZ was injected at the beginning of the experiment. Then the mice were received oral FMT inoculations (0.1 ml) once daily for two weeks (five weeks of age to seven weeks of age). Then the mice were regularly maintained (on respective diet) for another three weeks (ten weeks of age). Then, the mice were humanely euthanized to collect samples for different analyses (Fig. 1a; Study scheme).

**Fig. 1.**
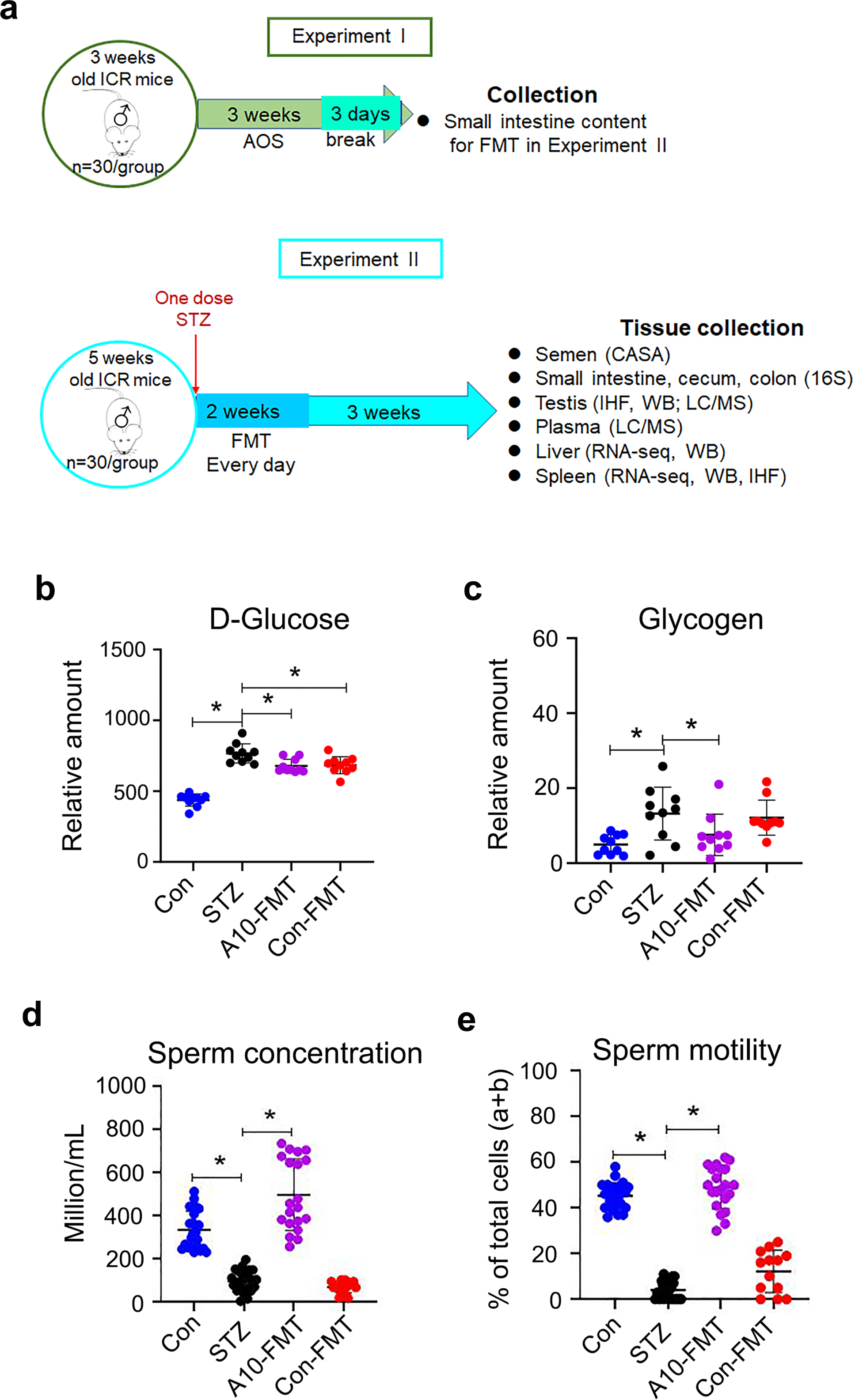
A10-FMT decreased blood glucose, and improved semen quality in type 1 diabetes. **a** Blood glucose levels. The y-axis represents the concentration (mmol/L). The x-axis represents the treatment (n = 30/group). \**p <* 0.05. **b** Blood glycogen levels. The y-axis represents the relative amount. The x-axis represents the treatment (n = 30/group). \**p <* 0.05. **c** Sperm concentration. The y-axis represents the concentration. The x-axis represents the treatment (n = 30/group). \**p <* 0.05. **d** Sperm motility. The y-axis represents the percentage of cells. The x-axis represents the treatment (n = 30/group). \**p <* 0.05.

### Evaluation of spermatozoa motility using a computer-assisted sperm analysis system

Spermatozoa motility was assessed using a computer-assisted sperm assay (CASA) method according to World Health Organization guidelines^22^. After euthanasia, spermatozoa were collected from the cauda epididymis of mice and suspended in DMEM/F12 medium with 10% FBS and incubated at 37.5 ℃ for 30 min; samples were then placed in a pre-warmed counting chamber. The micropic sperm class analyzer (CASA system) was used in this investigation. It was equipped with a 20-fold objective, a camera adaptor (Eclipse E200, Nikon, Japan), and a camera (acA780-75gc, Basler, Germany), and it was operated by an SCA sperm class analyzer (MICROPTIC S.L.). The classification of sperm motility was as follows: grade A linear velocity >22 μm s^-1^; grade B <22 μm s^-1^ and curvilinear velocity >5 μm s^-1^; grade C curvilinear velocity <5 μm s^-1^; and grade D = immotile spermatozoa. The spermatozoa motility data represented only grade A + grade B since only these two grades are considered to be functional.

#### Morphological observations of spermatozoa

*T*he extracted murine caudal epididymides were placed in RPMI medium, finely chopped, and then Eosin Y (1%) was added for staining as described previously^22^. Spermatozoon abnormalities were then viewed using an optical microscope and were classified into head or tail morphological abnormalities: two heads, two tails, blunt hooks, and short tails. The examinations were repeated three times, and 500 spermatozoa per animal were scored.

#### Assessment of acrosome integrity

After harvest, mouse spermatozoa were incubated at 37.5 ℃ for 30 min, after which a drop of sperm suspension was uniformly smeared on a clean glass slide. Smeared slides were air dried and incubated in methanol for 2 min for fixation. After fixation, the slides were washed with PBS three times. Assessment of an intact acrosome was accomplished by staining the sperm with 0.025% Coomassie brilliant blue G-250 in 40% methanol for 20 min at room temperature (RT). The slides were then washed three times with PBS and mounted with 50% glycerol in PBS. Acrosomal integrity was determined by an intense staining on the anterior region of the sperm head under bright-field microscopy (AH3-RFCA, Olympus, Tokyo, Japan) and scored accordingly^22^.

### RNA Isolation and RNA-seq analyses^22^

Briefly, total RNA was isolated using TRIzol Reagent (Invitrogen) and purified using a Pure-Link1 RNA Mini Kit (Cat: 12183018A; Life Technologies) following the manufacturers’ protocol. Total RNA samples were first treated with DNase I to degrade any possible DNA contamination. Then the mRNA was enriched using oligo(dT) magnetic beads. Mixed with the fragmentation buffer, the mRNA was broken into short fragments (about 200 bp), after which, the first strand of cDNA was synthesized using a random hexamer-primer. Buffer, dNTPs, RNase H, and DNA polymerase I were added to synthesize the second strand. The double strand cDNA was purified with magnetic beads. Subsequently, 3’-end single nucleotide A (adenine) addition was performed. Finally, sequencing adaptors were ligated to the fragments. The fragments were enriched by PCR amplification. During the QC step, an Agilent 2100 Bioanaylzer and ABI StepOnePlus Real-Time PCR System were used to qualify and quantify the sample library. The library products were prepared for sequencing in an Illumina HiSeqTM 2500. The reads were mapped to reference genes using SOAPaligner (v. 2.20) with a maximum of two nucleotide mismatches allowed at the parameters of “-m 0 -x 1000 - s 40 -l 35 -v 3 -r 2”. The read number of each gene was transformed into RPKM (reads per kilo bases per million reads), and then differentially expressed genes were identified using the DEGseq package and the MARS (MA-plot-based method with random sampling model) method. The threshold was set as FDR ≤0.001 and an absolute value of log_2_ ratio ≥1 to judge the significance of the difference in gene expression. Then on the data were analyzed by GO enrichment, KEGG enrichment.

### Sequencing of microbiota from intestine digesta samples and data analysis^22^

#### DNA Extraction

Total genomic DNA of small intestine, cecum and colon digesta was isolated using an E.Z.N.A.R Stool DNA Kit (Omega Bio-tek Inc., USA) following the manufacturer’s instructions. DNA quantity and quality were analyzed using NanoDrop 2000 (Thermo Scientific, USA) and 1% agarose gel. Ten samples/groups were determined.

#### Library preparation and sequencing

The V3-V4 region of the 16S rRNA gene was amplified using the primers MPRK341F (50-ACTCCTACGGGAGGCAGCAG -30) and MPRK806R: (50-GGACTACHVGGGTWTCTAAT -30) with Barcode. The PCR reactions (total 30 μL) included 15 μL PhusionR High-Fidelity PCR Master Mix (New England Biolabs), 0.2 mM primers, and 10 ng DNA. The thermal cycle was carried out with an initial denaturation at 98 ℃, followed by 30 cycles of 98 ℃ for 10 s, 50 ℃ for 30 s, 72 ℃ for 30 s, and a final extension at 72 ℃ for 5 min. PCR products were purified using a GeneJET Gel Extraction Kit (Thermo Scientific, USA). The sequencing libraries were constructed with NEB NextR UltraTM DNA Library Prep Kit for Illumina (NEB, United States) following the manufacturer’s instructions and index codes were added. Then, the library was sequenced on the Illumina HiSeq 2500 platform and 300 bp paired-end reads were generated at the Novo gene. The paired-end reads were merged using FLASH (V1.2.71). The quality of the tags was controlled in QIIME (V1.7.02), meanwhile all chimeras were removed. The “Core Set” of the Greengenes database3 was used for classification, and sequences with >97% similarity were assigned to the same operational taxonomic units (OTUs).

#### Analysis of sequencing data

Operational taxonomic unit abundance information was normalized using a standard of sequence number corresponding to the sample with the least sequences. The alpha diversity index was calculated with QIIME (Version 1.7.0). The Unifrac distance was obtained using QIIME (Version 1.7.0), and PCoA (principal coordinate analysis) was performed using R software (Version 2.15.3). The linear discriminate analysis effect size (LEfSe) was performed to determine differences in abundance; the threshold LDA score was 4.0. GraphPad Prism7 software was used to produce the graphs.

### Plasma and testis metabolite measurements by LC-MS/MS

Plasma samples were collected and immediately stored at -80 °C. Before LC-MS/MS analysis, the samples were thawed on ice and processed to remove proteins. Testis samples were collected and the same amount of tissue from each mouse testis was used to isolate the metabolites using CH3OH: H2O (V: V) = 4:1. Then samples were detected by ACQUITY UPLC and AB Sciex Triple TOF 5600 (LC/MS) as reported previously^17, 22^. Ten samples/groups were analyzed for plasma or testis samples. The HPLC conditions employed an ACQUITY UPLC BEH C18 column (100 mm × 2.1 mm, 1.7 μm), solvent A [aqueous solution with 0.1% (v/v) formic acid], and solvent B [acetonitrile with 0.1% (v/v) formic acid] with a gradient program. The flow rate was 0.4 mL/min and the injection volume was 5 μL. Progenesis QI v2.3 (Nonlinear Dynamics, Newcastle, UK) was implemented to normalize the peaks. Then the Human Metabolome Database (HMDB), Lipidmaps (v2.3), and METLIN software were used to qualify the data. Moreover, the data were processed with SIMCA software (version 14.0, Umetrics, Umeå, Sweden) following pathway enrichment analysis using the KEGG database (http://www.genome.jp/KEGG/pathway.html).

### Determination of blood glucose, insulin, ALT, TG, TC, T-AOC, GSH, SOD and catalase

Blood insulin was determined by the kit form Beijing Solarbio Science & Technology Co., Ltd (Beijing, P. R. China; Cat. #: SEKM0141). Blood glucose, ALT, TG, TC, T-AOC, SOD, and catalase were determined by the kits from Nanjing Jiancheng Bioengineering Institute [Nanjing, P.R. China; glucose (Cat. #: F006-1-1); ALT (Cat. #: C009-2-1); TG (Cat. #: A110-1-1); TC (Cat. #: A111-1-1); T-AOC (Cat. #: A015-2-1); GSH (Cat. #: A006-2-1); SOD (Cat. #: A001-3-2); Catalase (Cat. #: A007-1-1)] ^25^. All the procedures were followed from the manufacturer’s instructions.

### Measurement of iron content in spleen

The amount of ferric iron in the spleens was determined by Perl’s Prussian blue stain as described by Kohyama et al^26^. Spleen tissues were fixed with 4% paraformaldehyde, then embedded in paraffin. And 5 μm sections were cut and stained with Perl’s Prussian blue and pararosaniline (Sigma).

### Histopathological analysis

Testicular tissues were fixed in 10% neutral buffered formalin, paraffin embedded, cut into 5 μm sections and subsequently stained with hematoxylin and eosin (H&E) for histopathological analysis.

### Western blotting

Western blotting analysis of proteins was carried out as previously reported^17, 22^. Briefly, testicular tissue samples were lysed in RIPA buffer containing the protease inhibitor cocktail from Sangong Biotech, Ltd. (Shanghai, China). Protein concentration was determined using a BCA kit (Beyotime Institute of Biotechnology, Shanghai, China). Goat anti-actin was used as a loading control. The information for primary antibodies (Abs) were listed in Supplementary Table 1. Secondary donkey anti-goat Ab (Cat no.: A0181) was purchased from Beyotime Institute of Biotechnology, and goat anti-rabbit (Cat no.: A24531) Abs were bought from Novex^®^ by Life Technologies (USA). Fifty micrograms of total protein per sample were loaded onto 10% SDS polyacrylamide electrophoresis gels. The gels were transferred to a polyvinylidene fluoride (PVDF) membrane at 300 mA for 2.5 h at 4 ℃. The membranes were then blocked with 5% bovine serum albumin (BSA) for 1 h at RT, followed by three washes with 0.1% Tween-20 in TBS (TBST). The membranes were incubated with primary Abs diluted with 1:500 in TBST with 1% BSA overnight at 4 ℃. After three washes with TBST, the blots were incubated with the HRP-labelled secondary goat anti-rabbit or donkey anti-goat Ab respectively for 1 h at RT. After three washes, the blots were imaged. The bands were quantified using Image-J software. The intensity of the specific protein band was normalized to actin first, then the data were normalized to the control. The experiment was repeated >6 times.

### Detection of protein levels and location in testis using immunofluorescence staining

The methodology for immunofluorescence staining of testicular samples is reported in our recent publications^17, 22^. Sections of testicular tissue (5 μm) were prepared and subjected to antigen retrieval and immunostaining as previously described^17, 22^. Briefly, sections were first blocked with normal goat serum in PBS, followed by incubation with primary Abs (Supplementary Table 1; 1:100 in PBS-0.5% Triton X-100; Bioss Co. Ltd. Beijing, PR China) at 4 °C overnight. After a brief wash, sections were incubated with an Alexa 546-labeled goat anti-rabbit secondary Ab (1:100 in PBS; Molecular Probes, Eugene, OR, USA) at RT for 30 min and then counterstained with 4’,6-diamidino-2-phenylindole (DAPI). The stained sections were examined using a Leica Laser Scanning Confocal Microscope (LEICA TCS SP5 II, Germany). Ten animal samples from each treatment group were analysed. Positively stained cells were counted. A minimum of 1000 cells were counted for each sample of each experiment. The data were then normalized to the control.

### Statistical analysis

Data were analyzed using SPSS statistical software (IBM Co., NY) with one-way analysis of variance (ANOVA) followed by LSD multiple comparison tests or T-test. The data were shown as the mean ± SEM. Statistical significance was based on p < 0.05.

### Data availability

Liver and spleen RNA-seq raw data were deposited in NCBI’s Gene Expression Omnibus under accession number GSE184021 and GSE184023, respectively. The microbiota raw sequencing data generated in this study has been uploaded to the NCBI SRA database with the accession number PRJNA759114 (small intestine), PRJNA759089 (cecum), and PRJNA759063 (colon).

## Results

### A10-FMT decreased blood glucose, ameliorated STZ-induced T1D diminished semen quality, and impaired gut microbiota

Three days after STZ treatment (one dose, 135 mg/kg body weight), blood glucose was significantly higher in the STZ group [23.8 ± 2.1 mmol/L (mM)] than that in the control group (Con; 8.2 ± 1.3 mM) which indicated that the animals were diabetic (T1D; Fig. 1a; Study scheme). After five weeks treatment, the body weight and blood insulin were lower in STZ group than that in Con, while FMT from AOS improved gut microbiota (A10-FMT) produced a slight increase in body weight and blood insulin over STZ [no significant difference between STZ, A10-FMT, and FMT from gut microbiota of control animals (Con-FMT); Supplementary Fig. 1a and b]. After five weeks of treatment, blood glucose was significantly higher in STZ while it was significantly decreased by A10-FMT and Con-FMT (Fig. 1b) which indicated that FMT treatment improved T1D status^8^. At the same time blood glycogen was higher in STZ animals while it was reduced by A10-FMT but not Con-FMT (Fig. 1c), which further suggested that A10-FMT improved T1D status. STZ significantly diminished semen quality through decreasing sperm motility and concentration (Fig. 1d and e). However, A10-FMT significantly increased sperm motility and concentration, while Con-FMT produced a slight change (Fig. 1d and e). The data suggest that A10-FMT contained beneficial microbiota for improving semen quality. Gut dysbiosis has been reported in both T1D humans and animal models^9–11^.

Moreover, dysbiosis may also impair spermatogenesis to decrease semen quality. In the current investigation, we found a similar phenomenon as the microbiota in the small intestine, cecum, and colon were changed in T1D animals with an increase in the harmful bacteria *Bacteroides*, *Mycoplasma,* and *Escherichia*^8, 12^ (Fig. 2; Fig. 3; Supplementary Fig. 1c-n). A10-FMT increased the beneficial microbiota *Lactobacillus* in the small intestine, while decreasing the harmful bacteria *Escherichia* in the cecum and colon (Fig. 2d and h; Fig. 3d; Supplementary Fig. 1c-n). However, Con-FMT decreased *Bacteroides*, *Mycoplasma,* and *Escherichia,* but did not increase *Lactobacillus* (Fig. 2; Fig. 3; Supplementary Fig. 1c-n).

**Fig. 2.**
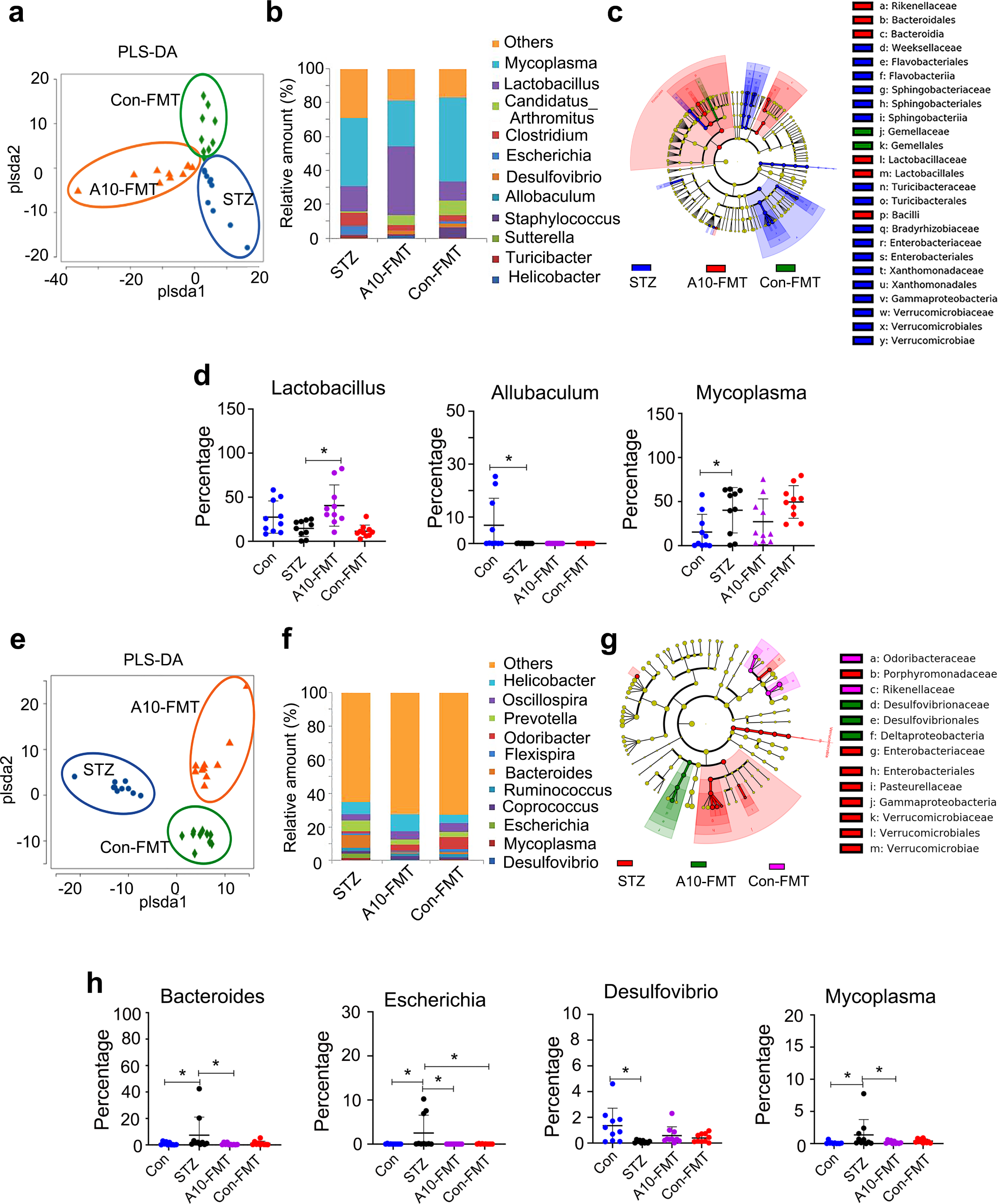
A10-FMT improved small intestinal and cecal microbiota in type 1 diabetes. **a** PLS-DA (OTU) of small intestine microbiota in HFD, A10-FMT, and Con-FMT groups. **b** Small intestine microbiota levels at the genus level in STZ, A10-FMT, and Con-FMT groups. The y-axis represents the relative amount (%). The x-axis represents the treatments. Different colors represent different microbiota. **c** Cladogram of the linear discriminate analysis effect size (LEfSe) determining the difference in abundance of small intestine microbiota. **d** Changed microbiota in the small intestine. The y-axis represents the relative amount at the genus level. The x-axis represents the treatment. \**p <* 0.05. **e** PLS-DA (OTU) of cecum microbiota in STZ, A10-FMT, and Con-FMT groups. **f** Cecum microbiota levels at the genus level in STZ, A10-FMT, and Con-FMT groups. The y-axis represents the relative amount (%). The x-axis represents the treatments. Different colors represent different microbiota. **g** Cladogram of the LEfSe determining the cecum microbiota difference in abundance. **h** Changed microbiota in the cecum. The y-axis represents the relative amount at the genus level. The x-axis represents the treatment. \**p <* 0.05.

**Fig. 3.**
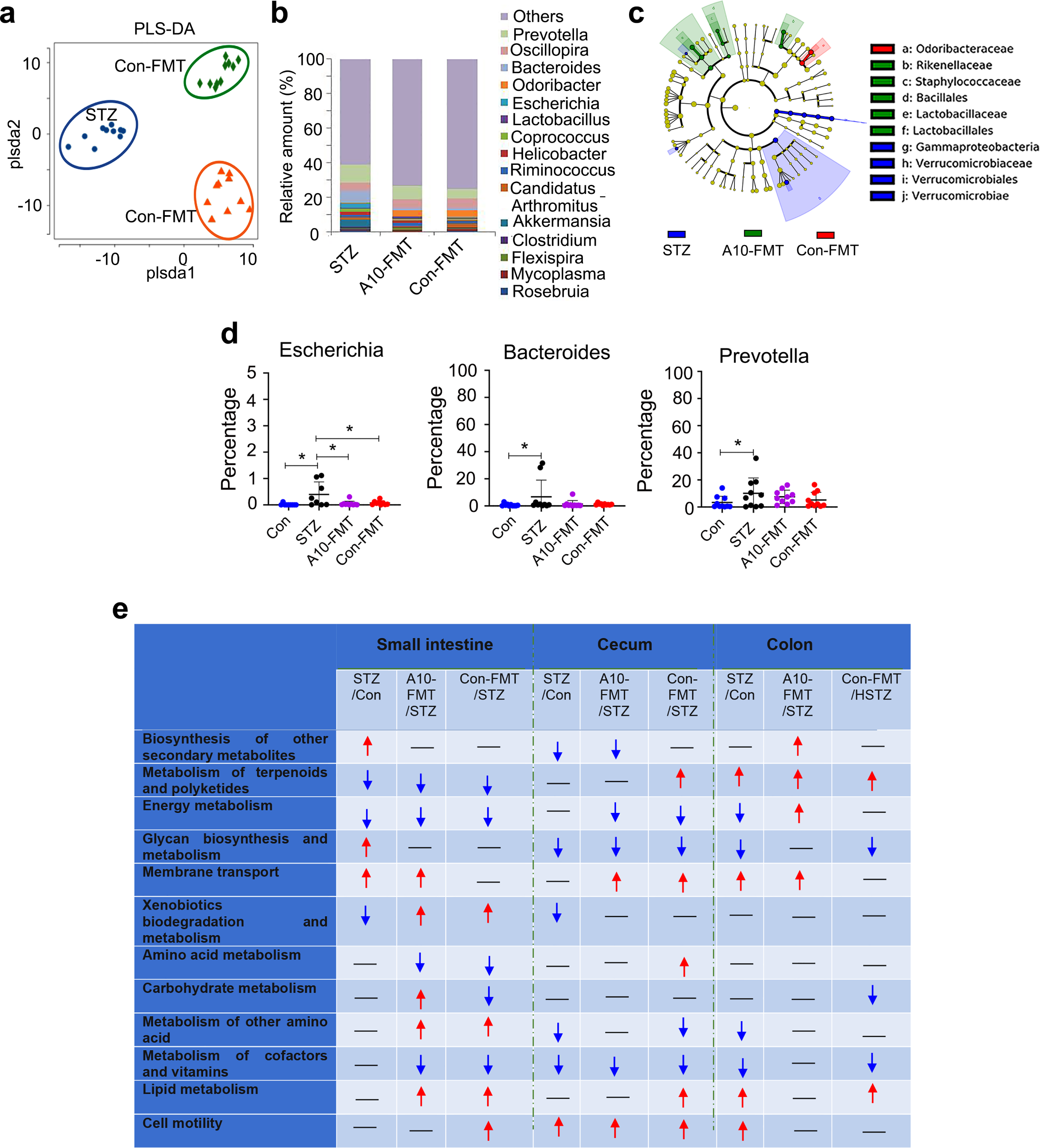
A10-FMT improved colon microbiota in type 1 diabetes. **a** PLS-DA (OTU) of colon microbiota in STZ, A10-FMT, and Con-FMT groups. **b** Colon microbiota levels at the genus level in STZ, A10-FMT, and Con-FMT groups. The y-axis represents the relative amount (%). The x-axis represents the treatments. Different colors represent different microbiota. **c** Cladogram of the LEfSe determining the difference in abundance of colon microbiota. **d** Changed microbiota in the colon. The y-axis represents the relative amount at the genus level. The x-axis represents the treatment. \**p <* 0.05. **e** Summary of signaling pathways of changed microbiota genes by Kyoto Encyclopedia of Genes and Genomes (KEGG) enrichment analysis. Red arrow indicates increased microbiota genes in each comparison. Blue arrow indicates decreased microbiota genes in each comparison.

The gut microbiota participates in host metabolism by interacting with host signaling pathways. Kyoto Encyclopedia of Genes and Genomes (KEGG) analysis of changed microbiota genes found that 12 major signaling pathways were upset by STZ and recovered by A10-FMT and/or Con-FMT in the colon, cecum, and/or small intestine (Fig. 3e). Interestingly, the “biosynthesis of other secondary metabolites” pathway was increased by A10-FMT specifically in the colon, and the “energy metabolism” pathway was decreased by STZ while increased by A10-FMT in the colon; meanwhile, the “carbohydrate metabolism” pathway was increased by A10-FMT while decreased by Con-FMT in the small intestine (Fig. 3e). However, the “metabolism of terpenoids and polyketides”, “amino acid metabolism”, and “lipid metabolism” pathways were only increased by Con-FMT in the cecum; the “carbohydrate metabolism” pathway was decreased only by Con-FMT in the colon; and the “cell motility” pathway was only increased by Con-FMT in the small intestine (Fig. 3e). Moreover, the “Energy metabolism” pathway in the cecum, “amino acid metabolism” pathway in the small intestine, and “metabolism of cofactor and vitamins” pathway in the small intestine were decreased by both A10-FMT and Con-FMT, while they were not changed by STZ; “membrane transport” pathway in the cecum, and “metabolism of other amino acids” and “lipid metabolism” in the small intestine were increased by both A10-FMT and Con-FMT while they remained unchanged by STZ; “xenobiotics biodegradation and metabolism” was decreased by STZ while increased by both A10-FMT and Con-FMT (Fig. 3e). The data indicated that A10-FMT and Con-FMT differentially modified gut microbiota and microbial function to affect blood metabolites and other processes such as spermatogenesis.

### A10-FMT-recovered gut microbiota improved the blood metabolome

T1D is a metabolic related disease, and gut microbiota influence the blood metabolome; therefore, we next explored blood metabolome changes using LC/MS (Data Set 1). KEGG enrichment analysis of changed microbiota genes indicated that the “carbohydrate metabolism” pathway was increased by A10-FMT in the small intestine, and it was interesting to note that blood carbohydrate was increased by STZ while decreased by A10-FMT or Con-FMT (Fig. 4a).

**Fig. 4.**
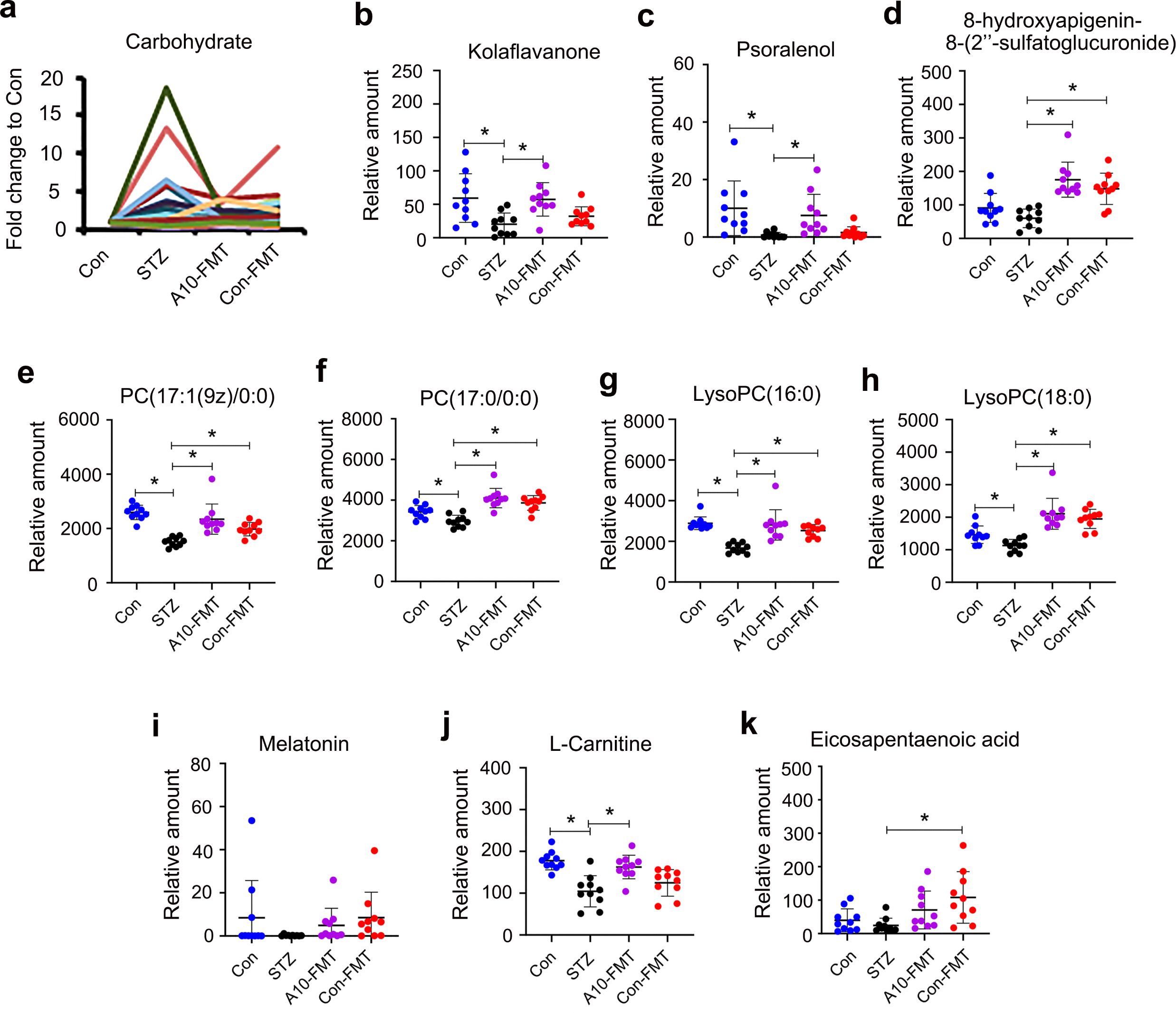
A10-FMT improved blood metabolism. **a** Blood carbohydrate levels in different treatments. The y-axis represents the fold change compared to control group (Con). The x-axis represents the treatment. **b** Blood flavonoid Kola flavanone levels in different treatments. The y-axis represents the relative amount. The x-axis represents the treatment. \**p <* 0.05. **c** Blood flavonoid Psoralenol levels in different treatments. The y-axis represents the relative amount. The x-axis represents the treatment. \**p <* 0.05. **d** Blood flavonoid 8-hydroxyaplgenin 8-(2’’-sulfatoglucuronide) levels in different treatments. The y-axis represents the relative amount. The x-axis represents the treatment. \**p <* 0.05. **e** Blood PC (17:1(9z)/0:0) levels in different treatments. The y-axis represents the relative amount. The x-axis represents the treatment. \**p <* 0.05. **f** Blood PC (17:0/0:0) levels in different treatments. The y-axis represents the relative amount. The x-axis represents the treatment. \**p <* 0.05. **g** Blood LysoPC (16:0) levels in different treatments. The y-axis represents the relative amount. The x-axis represents the treatment. \**p <* 0.05. **h** Blood LysoPC (18:0) levels in different treatments. The y-axis represents the relative amount. The x-axis represents the treatment. \**p <* 0.05. **i** Blood melatonin levels in different treatments. The y-axis represents the relative amount. The x-axis represents the treatment. \**p <* 0.05. **j** Blood L-Carnitine levels in different treatments. The y-axis represents the relative amount. The x-axis represents the treatment. \**p <* 0.05. **k** Blood eicosapentaenoic acid (EPA) levels in different treatments. The y-axis represents the relative amount. The x-axis represents the treatment. \**p <* 0.05.

Two other large clusters of compounds were decreased by STZ while increased by A10-FMT including flavonoids (Fig. 4b-d; Supplementary Fig. 2a), and glycerophosphocholines/ glycerophosphoethanolamines (Fig. 4e-h; Supplementary Fig. S2b-i). Flavonoids are compounds that play important roles in antioxidant activity and other functions to protect organisms. Glycerophosphocholines and glycerophosphoethanolamines have protective functions within the body.

In addition, melatonin, that has antioxidant effects along with many other functions, was decreased by STZ while increased by A10-FMT and Con-FMT (Fig. 4i) although at non-significant levels, which suggested that FMT could help increase systemic antioxidant capabilities. L-carnitine, an important compound involved in male sperm formation and function, was decreased by STZ while increased by A10-FMT but not by Co-FMT (Fig. 4j) which suggested that A10-FMT and Con-FMT differentially influence blood metabolism especially for antioxidant compound production. The data were consistent with gut microbiota data that A10-FMT increased *Lactobacillus* which have been shown to produce metabolites that improve liver or cardiac impairment^27–29^ (Fig. 3e). Most interestingly, blood n-3 polyunsaturated acid (PUFA) eicosapentaenoic acid (EPA) was decreased by STZ while increased by A10-FMT and Con-FMT (Fig. 4k). It is known that n-3 PUFA especially EPA and DHA are important for many aspects of our health, including spermatogenesis^30–32^.

### A10-FMT-improved blood metabolite ameliorated testicular metabolome (PUFA and retinoic acid) and the testicular microenvironment

The blood metabolome was upset in T1D (by STZ) while it recovered under treatment with A10-FMT and/or Con-FMT, which suggested that FMT creates beneficial systemic improvements in animals. Blood metabolites are important for testis growth and sperm development^17, 18, 22^; the blood metabolome and testicular metabolome are connected together, therefore, we set out to determine the testicular metabolome using LC/MS.

It was notable that the testicular n-3 PUFAs DHA and EPA were increased by A10-FMT but not by Con-FMT (Fig. 5a-c; Supplementary Fig. 3a; Data Set 2), which further suggested that A10-FMT is beneficial for testicular metabolism. The retinoic acid pathway plays a crucial role in spermatogenesis^33, 34^; and PUFA and retinoic acid signaling interact together to regulate spermatogenesis^35, 36^. STZ decreased retinol or retinoic acid related compounds while A10-FMT increased them (Fig. 5d and e; Supplementary Fig. 3b-d) which suggested that spermatogenesis was initiated by A10-FMT since retinoic acid turns meiosis on. Moreover, the protein levels of the spermatogonia cell marker genes PLZF and DAZL were elevated by A10-FMT (Fig. 5f) which confirmed initiation of the spermatogenesis process. Moreover, testosterone levels were increased by A10-FMT (Fig. 5g-i; Supplementary Fig. 3e-g); steroids other than testosterone were increased by A10-FMT while they were decreased by STZ (Fig. 5j and k; Supplementary Fig. 3h-l) which together further confirmed initiation of spermatogenesis. Similarly, in the blood, testicular glycerophosphocholines were increased by A10-FMT while they were decreased by STZ (Fig. 5l; Supplementary Fig. 3m-o). Melatonin metabolite 6-hydroxymelatonin, an active form of melatonin^37^, was increased by A10-FMT (Fig. 5m) which indicated that A10-FMT modulated both the systemic antioxidant status and also the testicular microenvironment to benefit spermatogenesis.

**Fig. 5.**
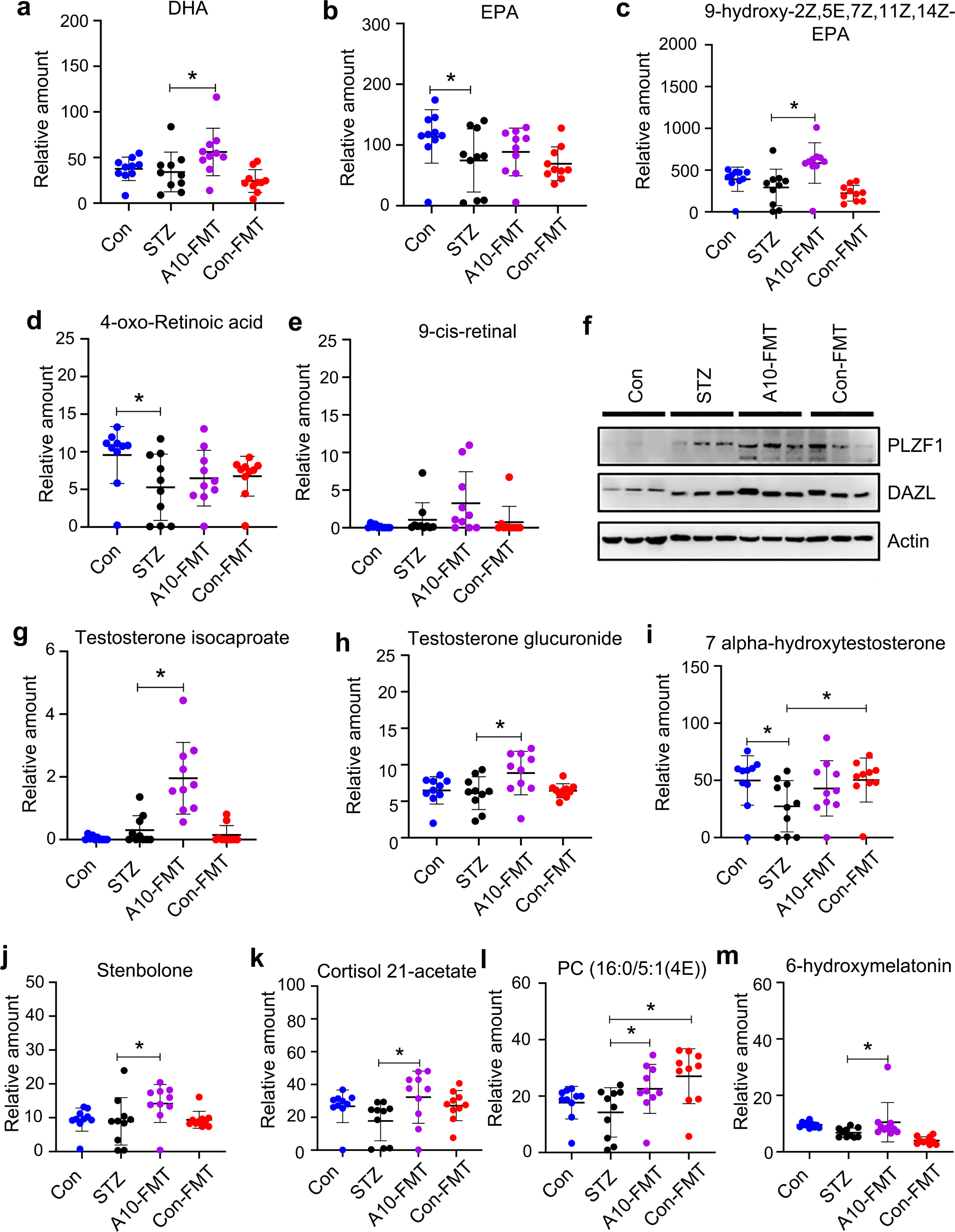
A10-FMT improved testicular metabolism. **a** Testicular docosahexaenoic acid (DHA) levels in different treatments. The y-axis represents the relative amount. The x-axis represents the treatment. **b** Testicular EPA in different treatments. The y-axis represents the relative amount. The x-axis represents the treatment. \**p <* 0.05. **c** Testicular 9-hydroxy-2Z,5E,7Z,11Z,14Z-Eicosapentaenoic acid levels in different treatments. The y-axis represents the relative amount. The x-axis represents the treatment. \**p <* 0.05. **d** Testicular 4-oxo-Retinoic acid levels in different treatments. The y-axis represents the relative amount. The x-axis represents the treatment. \**p <* 0.05. **e** Testicular 9-cis-retinal levels in different treatments. The y-axis represents the relative amount. The x-axis represents the treatment. \**p <* 0.05. **f** Testicular protein levels of *PLZF* and *DAZL* in different treatments determined by Western blotting. **g** Testicular testosterone isocaproate levels in different treatments. The y-axis represents the relative amount. The x-axis represents the treatment. \**p <* 0.05. **h** Testicular testosterone glucuronide levels in different treatments. The y-axis represents the relative amount. The x-axis represents the treatment. \**p <* 0.05. **i** Testicular 7alpha-hydroxytestosterone levels in different treatments. The y-axis represents the relative amount. The x-axis represents the treatment. \**p <* 0.05. **j** Testicular stenbolone levels in different treatments. The y-axis represents the relative amount. The x-axis represents the treatment. \**p <* 0.05. **k** Testicular cortisol 21-acetate levels in different treatments. The y-axis represents the relative amount. The x-axis represents the treatment. \**p <* 0.05. **l** Testicular PC (16:0/5:1(4E)) levels in different treatments. The y-axis represents the relative amount. The x-axis represents the treatment. \**p <* 0.05. **m** Testicular 6-hydroxymelatonin levels in different treatments. The y-axis represents the relative amount. The x-axis represents the treatment. \**p <* 0.05.

### A10-FMT-improved testicular microenvironment benefited spermatogenesis to improve semen quality

A10-FMT improved the testicular metabolome, especially through increased DHA and EPA which suggested that spermatogenesis should be improved. Spermatogenesis was indeed improved by A10-FMT (Fig. 6). The protein level (number of positive cells) of germ cell marker *VASA* (*DDX4*) was decreased in T1D animals while increased by A10-FMT (Fig. 6a and b). The protein level (number of positive cells) of the meiosis marker gene *SYCP3* was increased by A10-FMT while reduced by STZ (Fig. 6a and c). The protein level (number of positive cells) of transition protein 1 (*TP1*) was increased by A10-FMT (Fig. 6a and d). The protein level (number of positive cells) of the sperm protein *PGK2* was decreased by STZ while increased by A10-FMT (Fig. 6a and e). Moreover, the protein levels of some of the important genes for spermatogenesis *CREM*, *B-MYB*, *PIWIL1*, *ODF1*, *PGK2*, and *TP1* were determined by Western blotting. It was noteworthy that all these proteins were elevated by A10-FMT (Fig. 6f and g) which confirmed the IHF data and also suggested that spermatogenesis was improved by A10-FMT. At the same time the Sertoli cell marker gene *SOX9* was detected by IHF; it was shown that the number of *SOX9* positive cells remained unchanged by STZ or A10-FMT compared to Con (Fig. 6h). The data suggested that STZ impaired germ cells, but not somatic cells, to diminish spermatogenesis.

**Fig. 6.**
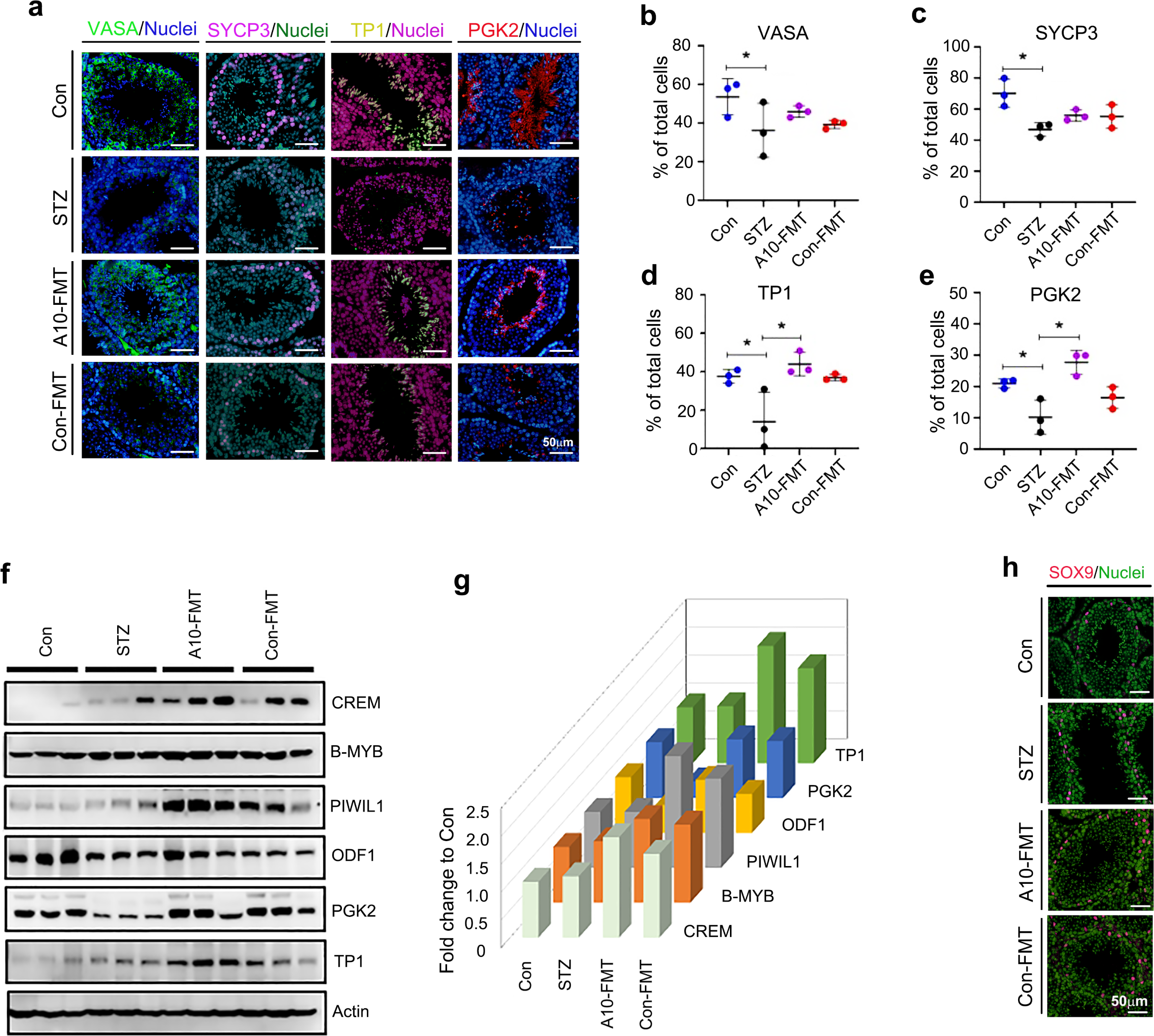
A10-FMT improved spermatogenesis process. **a** IHF staining of testicular germ cell marker *VASA*, meiosis marker *SYCP3*, sperm protein *PGK2*, and transition protein 1 (*TP1*) in each treatment. Scale bar: 50μm. **b** Quantitative data for *VASA* IHF staining. The y-axis represents the percentage of total cell. The x-axis represents the treatment. \**p <* 0.05. **c** Quantitative data for *SYCP3* IHF staining. The y-axis represents the percentage of total cell. The x-axis represents the treatment. \**p <* 0.05. **d** Quantitative data for *TP1* IHF staining. The y-axis represents the percentage of total cell. The x-axis represents the treatment. \**p <* 0.05. **e** Quantitative data for *PGK2* IHF staining. The y-axis represents the percentage of total cells. The x-axis represents the treatment. \**p <* 0.05. **f** Western blotting analysis of the proteins of important genes for spermatogenesis in each treatment. **g** Quantitative data for Western blotting analysis. **h** IHF staining of Sertoli cell marker *SOX9*.

### Furthermore, A10-FMT-improved gut microbiota benefited spleen immune function and liver function to strengthen the systemic environment for spermatogenesis

The spleen plays vital roles in systemic immune function^38, 39^. Recently, it has been established that the gut microbiota is involved in spleen development and function^38, 39^; and a gut-spleen interaction (axis) has been identified. In the current investigation, STZ also disrupted spleen function. Spleen RNA-seq data showed that STZ upset spleen gene expression while this was recovered by A10-FMT and/or Con-FMT (Supplementary Fig. S4). Gene enrichment analysis also showed that functions upset by STZ were reversed by A10-FMT or Con-FMT (Fig.7a; Supplementary Fig. 4a and b). “Adapted immune response”, “Complement cascade”, and “Platelet activation”, “Complement and coagulation cascade”, and “hemostasis” functional pathways were enriched for the genes decreased by STZ while they were increased by A10-FMT, which indicated that the immune and vasculature systems may be affected by STZ and recovered by A10-FMT. The “hemostasis” functional pathway indicated that STZ affected blood supply to the spleen, and it was confirmed by the expression of hemoglobin scavenger receptor (CD163) protein. CD163 is known to play major roles in the clearance and endocytosis of hemoglobin/haptoglobin complexes^26^. STZ decreased CD163 protein levels in the spleen while these were recovered by A10-FMT (Fig.7b).

**Fig. 7.**
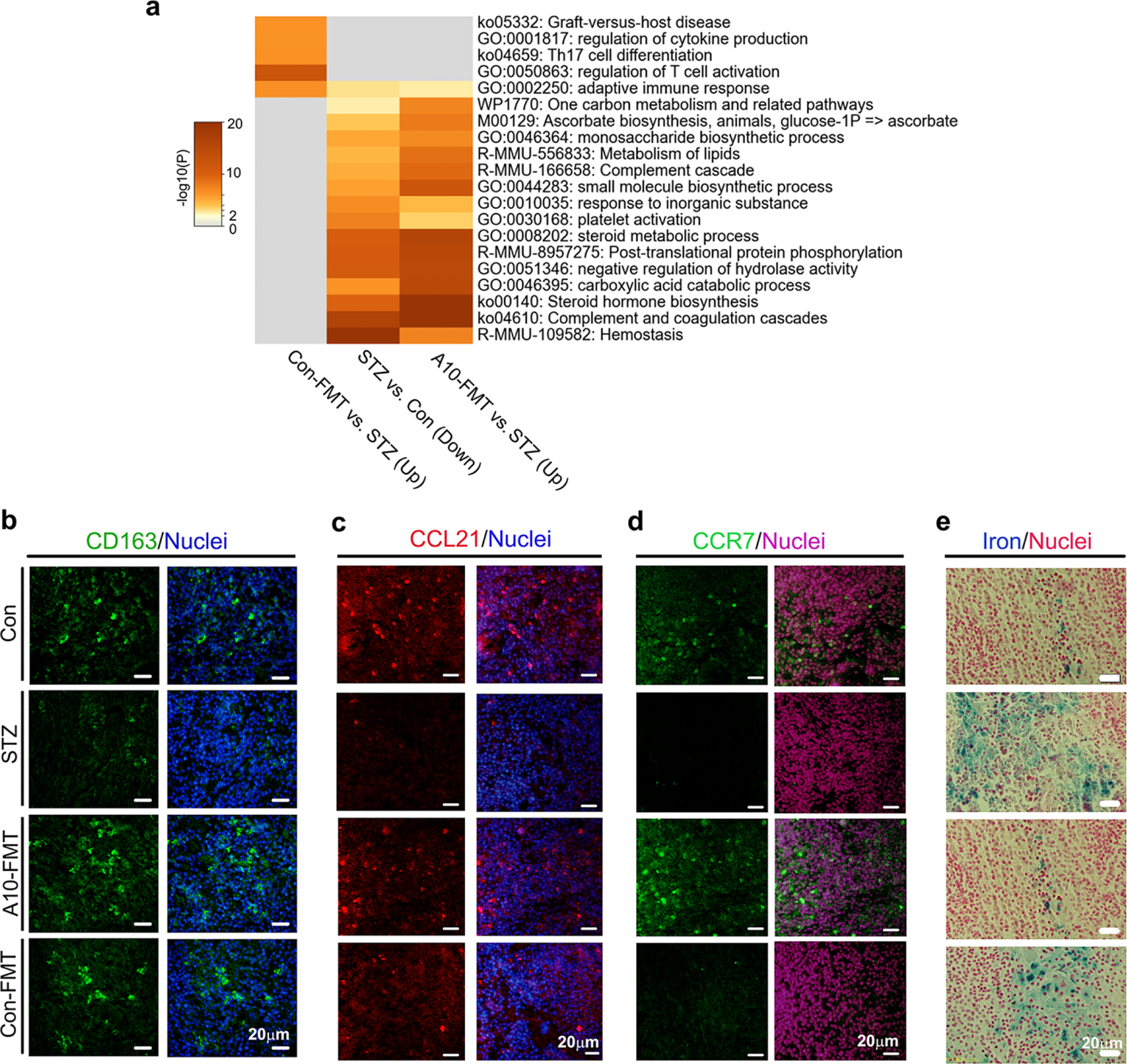
A10-FMT improved spleen function. **a** Functional enrichment analysis of STZ decreased genes while they were increased by A10-FMT or Con-FMT. **b** IHF staining of *CD163* in spleen tissue. **c** IHF staining of *CCL21* in spleen tissue. **d** IHF staining of *CCR7* in spleen tissue. **e** Perl’s Prussian blue stain for ferric iron in spleen tissue.

A10-FMT improved spleen immune functions. T-cell protein CCL21 and its receptor CCR7 play crucial roles in maintaining the active migratory state of T cells^40^. STZ decreased CCL21 protein levels (or positive cells) in the spleen while this was reversed by A10-FMT but not Con-FMT (Fig.7c). At the same time, the protein levels (or positive cells) of CCR7 were reduced by STZ while they were recovered by A10-FMT but not Con-FMT (Fig.7d). Moreover, the spleen plays important roles in iron homeostasis through the resorption of effete erythrocytes and the subsequent return of iron to the circulation. Free iron has the potential to become cytotoxic when electron exchange with oxygen is unrestricted and catalyzes the production of reactive oxygen species. Therefore, the balance of iron in the spleen and circulation is very important for health^26, 41, 42^. STZ increased the levels of iron in the spleen while this was reversed by A10-FMT but not Con-FMT (Fig.7e). Furthermore, the proliferation and apoptosis status of the spleen was recovered by A10-FMT (Supplementary Fig. 4c, d). The protein levels (or the number of positive cells) of cell proliferation marker Ki67 were reduced by STZ while they were increased by A10-FMT (Supplementary Fig. 4c). Furthermore, the protein levels of apoptosis markers p53 and Bax were diminished by STZ while they were recovered by A10-FMT (Supplementary Fig. 4d). All the data suggested that A10-FMT benefited spleen functions to assist with systemic immune functions since T1D is an important immune related disease^1^.

The liver plays a vital role in glucose metabolism and detoxification and many other functions maintain homeostasis. STZ upset liver function while this was recovered by A10-FMT (Supplementary Fig. 5). The liver damage marker alanine aminotransferase (ALT) was increased in the blood while this was decreased by A10-FMT (Supplementary Fig. 5a). RNA-seq analysis also showed that STZ disrupted liver functions while this was reversed by A10-FMT and/or Con-FMT (Supplementary Fig. 5b-d). KEGG enrichment analysis indicated that liver lipid metabolism was upset by STZ while it was reversed by A10-FMT and/or Con-FMT (Supplementary Fig. 5c and d). Furthermore, blood triglyceride (TG) and total cholesterol (TC) levels were increased by STZ while these were reversed by A10-FMT and/or Con-FMT (Supplementary Fig. 5e and f).

Antioxidants also play important roles in maintaining systemic functions. Blood total antioxidant capability (T-AOC) level was reduced by STZ while it was recovered by A10-FMT (Supplementary Fig. 5g). At the same time the levels of anti-oxidant enzymes SOD and catalase were decreased by STZ while increased by A10-FMT (Supplementary Fig. 5h and i). The antioxidant compound GSH was also increased by A10-FMT (Supplementary Fig. 5j). At the same time liver apoptosis status was upset by STZ while it was recovered by A10-FMT (Supplementary Fig. 5k). All the data suggested that A10-FMT had a strong capacity to improve levels of systemic antioxidants to benefit spermatogenesis since T1D induced hyperglycemia causes systemic oxidative stress^4^.

## Discussion

Since gut dysbiosis and T1D are closely related, and 90% of male T1D patients have varying degrees of testicular dysfunction and male infertility; gut dysbiosis and male infertility are also correlated. As most male T1D patients are of reproductive age, it is worth exploring protocols for improving spermatogenesis and male fertility^21^. Many studies have used different procedures to increase fertility, such as resveratrol and metformin, which are capable of improving semen quality to some extent^20, 21^. Very recently, we found that FMT from AOS-improved gut microbiota rescues high fat diet disrupted spermatogenesis. In the current investigation, we found that FMT from AOS-improved gut microbiota (A10-FMT), but not FMT from gut microbiota of control animals (Con-FMT), significantly increased sperm concentration and motility in STZ-induced T1D animals. Moreover, A10-FMT significantly decreased blood glucose and glycogen which suggested that A10-FMT may be supportive management for T1D patients.

Higher levels of gut *Bacteroidetes* have been found in T1D patients compared with their peers^12^. We found similar results; *Bacteroidetes* was increased in the cecum and colon of STZ-induced T1D animals compared with control group. Both A10-FMT and Con-FMT decreased the amount of *Bacteroidetes*. Meanwhile, *Prevotella* has been found to be inversely related to pancreatic beta cell function, and *Desulfovibrio* to be positively correlated with beta cell function^12^. In the current study, we found *Desulfovibrio* was decreased in the cecum of STZ-induced T1D animals while A10-FMT and Con-FMT restored its levels; *Prevotella* was increased in the colon of STZ-induced T1D animals while A10-FMT and Con-FMT decreased its levels. As these results show beneficial outcomes, this suggests that FMT could improve gut microbiota.

Of great interest in the current study was our finding that the small intestine *Lactobacillus* population, which has been shown to have multiple functions in human health^27–29, 43, 44^, and is discussed below, was increased by A10-FMT but not by Con-FMT. *Lactobacillus plantarum* 299v is reported to improve vascular endothelial function and decrease systemic inflammation in men with coronary artery disease and it is suggested that circulating gut-derived metabolites likely account for these improvements^27^. *Lactobacillus helveticus* R0052 has anti-inflammatory properties through downregulating Toll-like receptors, tumor necrosis factor-α, and nuclear factor-κb transcription in liver samples and decreasing proinflammatory cytokine plasma concentrations to alleviate hepatic injuries^28^. Meanwhile, *L. helveticus* R0052 is known to improve carbohydrate and fatty acid metabolism and reduce lithocholic acid levels^28^. Lew et al. report that some selected *Lactobacillus* strains improve lipid profiles via activation of energy and lipid metabolism, suggesting the potential of *Lactobacillus spp*. as promising natural interventions for the alleviation of cardiovascular and liver diseases^29^. *Lactobacillus rhamnosus* GG ATCC53103 and *Lactobacillus plantarum* JL01, can improve growth performance and immunity of piglets, and relieve weaning stress-related immune disorders such as intestinal infections and inflammation^43^. The enhanced immunity in the latter study took place through increasing levels of tauroursodeoxycholic acid (TCDA) and docosahexaenoic acid (DHA); however there was a simultaneous decrease in succinic and palmitic acids^43^. *Lactobacillus plantarum* PCA 236 has been shown to beneficially modulate goat fecal microbiota and milk fatty acid composition^44^. These studies indicate that *Lactobacillus spp.* have the capacity to modify the production of polyunsaturated fatty acids such as DHA in animal blood or other organs.

It is known that DHA is crucial for spermatogenesis and male reproductive functions^30–32^; it has also been shown that DHA supplementation can fully restore fertility and spermatogenesis in male mice^31^. Meanwhile, Gallardo et al. report that a high fat diet decreases testicular DHA levels, which may be related to the production of dysfunctional spermatozoa^32^. The identification of beneficial n-3 PUFAs in this study was enlightening; we suggest that DHA and EPA were increased by A10-FMT in the blood and testes which was correlated with the increase in *Lactobacillus* in the small intestine following A10-FMT treatment. The current and previous studies have demonstrated that *Lactobacillus spp.* have the potential to improve DHA levels, which is important for aspects of health, including improvements in spermatogenesis. Furthermore, testosterone is a strategic player in spermatogenesis, and approximately 94.4% of diabetes cases are associated with hypotestosteronemia. Moreover, the incidence of sexual and reproductive dysfunction in diabetic patients is 5–10-fold higher than that in nondiabetic individuals^21^. In current study, A10-FMT increased the testicular testosterone levels, which may account for the improvement of spermatogenesis and semen quality.

T1D is an immune related disease, and perturbations in the gut microbiota can impair the functions of immune cells and vice-versa^10^. Such dysbiosis is often detected in T1D subjects, especially those with an adverse immunoresponse^10^. As an organ of immunity, the spleen plays important roles in our health^38^; furthermore, it has been established that the gut-spleen axis affects spleen development and human health^38, 39^. In the current study, we found that STZ-induced T1D disrupted spleen function while A10-FMT rescued it through the improvement of immune cell function and iron levels. Free iron levels in the spleen or liver may potentially become cytotoxic as they can catalyze the production of reactive oxygen species (ROS)^41^. A10-FMT beneficially decreased spleen iron levels, reduced apoptosis protein levels, and increased cell proliferation marker ki67 levels, all of which indicate improvements in spleen function ^26^. Macrophages, indispensable immune cells, also play important roles in the regulation of spleen iron levels^41, 42^, which further suggests that A10-FMT improves spleen immune function. This improvement in spleen function may support systemic health and spermatogenesis.

The main characteristic of T1D is hyperglycemia which can induce oxidative stress, increase endoplasmic reticulum stress, and impair mitochondrial function^4^. We found that STZ-induced T1D decreased T-AOC level and the levels of some antioxidant enzymes, such as SOD, while A10-FMT restored them. We also detected that STZ-induced T1D caused liver damage through increasing ALT levels, while A10-FMT recovered it. At the same time A10-FMT improved liver function by modulating the apoptotic status in liver cells.

In summary, in our animal model, STZ-induced T1D disrupted spermatogenesis to diminish semen quality through decreasing sperm concentration and sperm motility. Most importantly, STZ-induced T1D caused gut dysbiosis. A10-FMT and Con-FMT decreased blood glucose levels and improved gut microbiota through the reduction of “harmful” microbes and increase in “beneficial” microbes. Most importantly, A10-FMT enhanced specific beneficial microbiota such *Lactobacillus* to increase the production of n-3 PUFA, such as DHA and EPA, to ameliorate spermatogenesis and semen quality while Con-FMT did not. Moreover, A10-FMT specifically improved spleen and liver function to promote sperm development and increase semen quality. Thus, AOS improved gut microbiota may support the improvement of semen quality and male fertility in T1D patients.

## Acknowledgements

We thank the investigators and staff of The Beijing Genomics Institute (BGI) and Shanghai LUMING Biotechnology CO., LCD for technical support. This study was supported by the National Natural Science Foundation of China (31772408 to YZ; 31672428 to HZ).

## Author contributions

Y.H., Y.F., X.Y., L.C., R.Z., T.M., B.Y., H.H., Y. Zhou, X.T., S.W., L.L., P.Z., and B.H. performed the experiments and analyzed the data. Y. Zhao., H.Z., W.S., Q.S., Z.S., and Y.R. designed and supervised the study. Y. Zhao. and H.Z. wrote the manuscript. All the authors edited the manuscript and approved the final manuscript.

## Competing interests

The authors declare no competing interests.

## Supplementary information

**Supplementary Fig. 1. Body weight and gut microbiota changes (STZ vs. Con)**. **a** Animal bodyweight. The y-axis represents the body weight (g). The x-axis represents the age (weeks). **b** Blood insulin levels. The y-axis represents the concentration (mIU/L). The x-axis represents the treatment. **c** The alpha index of the small intestine microbiota (Chao index). The y-axis represents the relative amount. The x-axis represents the treatment. **d** The beta index of small intestinal microbiota. The y-axis represents the relative amount. The x-axis represents the treatment. **e** PLS-DA (OTU) of small intestine microbiota in STZ and Con groups. **f** Small intestine microbiota levels at the genus level in STZ and Con groups. The y-axis represents the relative amount (%). The x-axis represents the individual microbiota. **g** The alpha index of the cecum microbiota (Chao index). The y-axis represents the relative amount. The x-axis represents the treatment. **h** The beta index of cecum microbiota. The y-axis represents the relative amount. The x-axis represents the treatment. **i** PLS-DA (OTU) of cecum microbiota in STZ and Con groups. **j** Cecum microbiota levels at the genus level in STZ and Con groups. The y-axis represents the relative amount (%). The x-axis represents the individual microbiota. **k** The alpha index of the colon microbiota (Chao index). The y-axis represents the relative amount. The x-axis represents the treatment. **l** The beta index of colon microbiota. The y-axis represents the relative amount. The x-axis represents the treatment. **m** PLS-DA (OTU) of colon microbiota in STZ and Con groups. **n** Colon microbiota levels at the genus level in STZ and Con groups. The y-axis represents the relative amount (%). The x-axis represents the individual microbiota.

**Supplementary Fig. 2. Blood metabolism changes**. **a** Blood flavonoid Malvidin 3-O-(6-O-(4-O-feruloyl-alpha-rhamnopyranosyl)-beta-glucopyranoside)-5-beta-glucopy-ranoside levels. The y-axis represents the relative amount (%). The x-axis represents the treatments. **b** Blood glycerophosphocholine levels in different treatments. The y-axis represents the fold change compared to control group (Con). The x-axis represents the treatment. **c** Blood LysoPC (15:0) levels. The y-axis represents the relative amount (%). The x-axis represents the treatments. **d** Blood LysoPC [18:1(9z)] levels. The y-axis represents the relative amount (%). The x-axis represents the treatments. **e** Blood LysoPC [22:6(4Z,7Z,10Z,13Z,16Z,19Z)] levels. The y-axis represents the relative amount (%). The x-axis represents the treatments. **f** Blood PE [22:2(13Z,16Z)/0:0] levels. The y-axis represents the relative amount (%). The x-axis represents the treatments. **g** Blood PE (22:0/0:0) levels. The y-axis represents the relative amount (%). The x-axis represents the treatments. **h** Blood PE [22:1(11Z)/0:0] levels. The y-axis represents the relative amount (%). The x-axis represents the treatments. **i** Blood LysoPE [0:0/24:6(6Z,9Z,12Z,15Z,18Z,21Z)] levels. The y-axis represents the relative amount (%). The x-axis represents the treatments.

**Supplementary Fig. 3. Testicular metabolite changes**. **a** Testicular 4,8,12,15,18-eicosapentaenoic acid levels. The y-axis represents the relative amount (%). The x-axis represents the treatments. **b** Testicular retinoids levels in different treatments. The y-axis represents the fold change compared to the control group (Con). The x-axis represents the treatment. **c** Testicular retinol levels. The y-axis represents the relative amount (%). The x-axis represents the treatments. **d** Testicular retinyl ester levels. The y-axis represents the relative amount (%). The x-axis represents the treatments. **e** Testicular testosterone levels in different treatments. The y-axis represents the fold change compared to control group (Con). The x-axis represents the treatment. **f** Testicular testosterone propionate levels. The y-axis represents the relative amount (%). The x-axis represents the treatments. **g** Testicular testosterone acetate levels. The y-axis represents the relative amount (%). The x-axis represents the treatments. **h** Testicular steroids levels in different treatments. The y-axis represents the fold change compared to control group (Con). The x-axis represents the treatment. **i** Testicular 3b,17b-Dihydroxyetiocholane levels. The y-axis represents the relative amount (%). The x-axis represents the treatments. **j** Testicular 5-Androstene-3b,16b,17a-triol levels. The y-axis represents the relative amount (%). The x-axis represents the treatments. **k** Testicular 3beta,17alpha,21-Trihydroxy-pregnenone levels. The y-axis represents the relative amount (%). The x-axis represents the treatments. **l** Testicular asterogenol levels. The y-axis represents the relative amount (%). The x-axis represents the treatments. **m** Testicular PCs levels in different treatments. The y-axis represents the fold change compared to the control group (Con). The x-axis represents the treatment. **n** Testicular) PC(4:0/18:1(9Z) level. The y-axis represents the relative amount (%). The x-axis represents the treatments. **o** Testicular PC (8:0/8:0) levels. The y-axis represents the relative amount (%). The x-axis represents the treatments.

**Supplementary Fig. 4. Additional data for spleen**. **a** PCA for RNA-seq analysis of spleen. **b** The functional enrichment analysis of STZ increased genes while these were decreased by A10-FMT or Con-FMT. **c** IHF staining of *ki67* in the spleen. **d** Western blotting analysis of *p53*, *Bax* and *Bcl-xl* in the spleen.

**Supplementary Fig. 5. A10-FMT improved liver function and systemic anti-oxidative capability**. **a** Blood alanine aminotransferase (ALT) levels. The y-axis represents the relative amount (%). The x-axis represents the treatments. **b** PCA for RNA-seq analysis of liver. **c** The functional enrichment analysis of STZ decreased genes while these were increased by A10-FMT or Con-FMT in the liver. **d** The functional enrichment analysis of STZ increased genes while these were decreased by 882 A10-FMT or Con-FMT in the liver. **e** Blood total triglyceride (TG) levels. The y-axis 883 represents the relative amount (%). The x-axis represents the treatments. **f** Blood total 884 cholesterol (TC) levels. The y-axis represents the relative amount (%). The x-axis 885 represents the treatments. **g** Blood total antioxidant capability (T-AOC) levels. The y-886 axis represents the relative amount (%). The x-axis represents the treatments. **h** Blood 887 total SOD levels. The y-axis represents the relative amount (%). The x-axis represents 888 the treatments. **i** Blood catalase levels. The y-axis represents the relative amount (%). 889 The x-axis represents the treatments. **j** Blood glutathione levels. The y-axis represents 890 the relative amount (%). The x-axis represents the treatments. **k** Western blotting 891 analysis of *Bax* and *Bcl-xl* in the liver.

**Supplementary Table 1. Primary antibody information.**

Data Set 1. Blood metabolites raw data.

Data Set 2. Testicular metabolites raw data.

